# Macrophage-mediated bystander T cell activation in varicose veins

**DOI:** 10.64898/2026.09.01.748721

**Authors:** Doğa Uncuer, Marie-Helen Winter, Tabea Wieler, Katja Buschmann, Bettina Kleis- Fischer, Carolin Mitschang, Sonja Leson, Matthias Klein, Tobias Bopp, Katrin Schäfer, Fatemeh Shahneh, Verena K. Raker, Christian Becker

**Affiliations:** Department of Dermatology and Allergology, University Hospital Augsburg, Germany; Department of Dermatology, University Medical Center Mainz, Johannes Gutenberg-University Mainz, Germany; Department of Cardiac and Vascular Surgery, University Medical Center Mainz, Johannes Gutenberg-University Mainz, Germany; Department of Dermatology, University Hospital Münster, Germany; Institute for Immunology, University Medical Center Mainz, Johannes Gutenberg-University Mainz, Germany; Department of Cardiology, University Medical Center Mainz, Johannes Gutenberg-University Mainz, Germany; Center for Thrombosis and Hemostasis, University Medical Center Mainz, Johannes Gutenberg-University Mainz, Germany

## Abstract

**Background:** Up to one-third of all adults are affected by chronic venous insufficiency, with varicose veins being a common manifestation of the condition. It is believed that inflammation plays a central role in the progression of the disease; however, the immune cell populations and mechanisms involved remain unclear.

**Methods:** Varicose veins were subjected to a stepwise mechanical and enzymatic dissociation, and the mononuclear cells resulting from these steps were characterized by flow cytometry. CD206-negative connective tissue macrophages and CD206-positive venous wall macrophages were sorted from five donors and subjected to transcriptomic profiling. Cytokine production by T cells in varicose veins was characterized following inhibition of intracellular protein transport. The ability of cytokines identified in macrophage transcriptomes to induce cytokine production in CD8^+^ and CD4^+^ effector and memory T cells was investigated using combinations of recombinant cytokines.

**Results:** Varicose veins harbor CD206^neg^ IL-1α^+^ S100A8/9^+^ macrophages in the connective tissue and pro-resolving and tissue regenerating CD206^+^ TIMD4^+^ LYVE1^+^ macrophages in the vein wall. T cells predominantly localize within the connective tissue, express an effector-memory T cell phenotype and produce IFN-γ in situ. Cytokines identified in the macrophage transcriptome collectively induced IFN-γ, as well as TNF-α, IL-22, and GM-CSF in effector memory T cells; in addition to IL-15 and IL-18, TNF-α and IL-10 proved to be key drivers.

**Conclusions:** The identified macrophage-T cell cytokine axis contributes to chronic venous insufficiency pathogenesis and represents a potential therapeutic target.

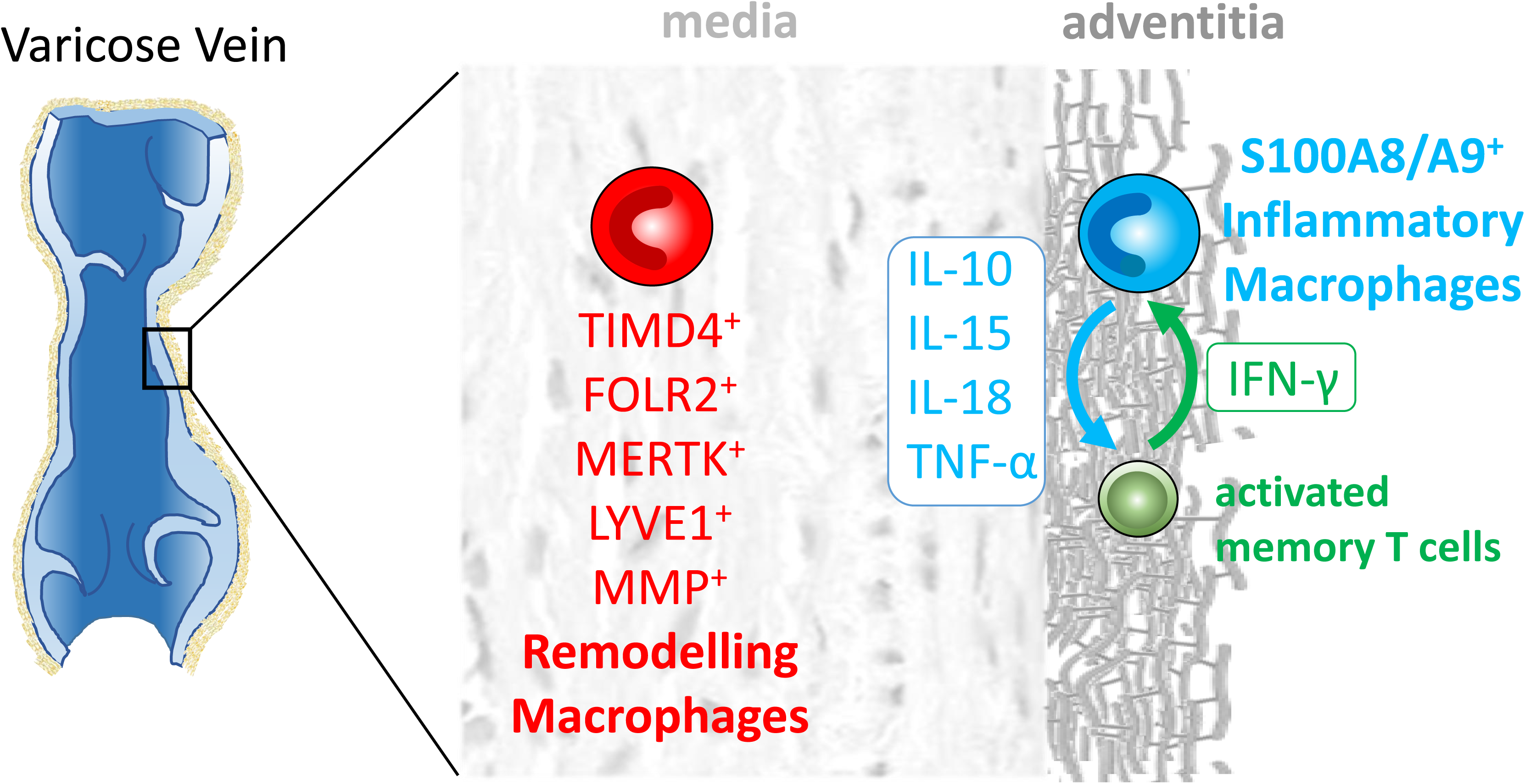

## Introduction

Chronic venous insufficiency (CVI) affects up to one-third of the adult population.^1^ Varicose veins are among its most common clinical manifestations. Inflammation is not merely a concomitant symptom but the primary driver of the disease’s progression to a chronic state.^2^ Accordingly, varicose veins exhibit a higher infiltration of immune cells than healthy veins^3^ with myeloid cells accounting for a significant proportion of this infiltrate.^3, 4^ Some work studies suggest a role of macrophage-released factors in CVI pathology^3–5^ but their functional state in varicose veins and possible interaction with other immune or stromal cells remain unclear. We have previously reported that varicose veins contain CD69^+^ tissue-resident effector-memory T cells.^6^ To characterize macrophages in varicose veins and their potential local interaction with T-cells, we performed stepwise mechanical purification/tissue digestion and expression profiling of myeloid cells sorted from varicose veins. We further investigated T-cell cytokine production in situ and the effect of macrophage cytokines, identified from the transcriptome of macrophages in varicose veins, on cytokine production by CD8^+^ and CD4^+^ effector-memory T cells.

## Methods

### Patient samples

Varicose vein samples were obtained from patients undergoing vein stripping for primary varicosity at the Departments of Dermatology in Mainz and Münster, Germany. Bypass vein samples were received from the Department of Cardiothoracic and Vascular Surgery, Mainz. In Mainz, tissue samples were used following voluntary ‘broad consent’ for the use of routinely collected material. In Münster, the study received approval by the ethics committee of the Westphalia-Lippe Medical Association (2025-363-f-S). All experimental procedures are in line with the guideline of the Declaration of Helsinki. Vein samples were treated anonymously. None of the samples contained blood clots.

### Stepwise mechanical processing and Tissue digestion

The surgically harvested venous tissue was processed immediately. First, the samples were cut lengthwise with scissors in a Petri dish, and any residual blood was removed. The tissue was then chopped into 1-mm pieces on ice using scalpels. The tissue pieces were washed with PBS and incubated in 5 mL of HBSS buffer containing 450 U/mL collagenase IV (Worthington Biochemical Corporation, CLS-4, 1 gm, CAT# LS004188), 500 U/mL DNase I (grade II, Roche, REF 10104159001), 1 mg/mL trypsin inhibitor (Worthington Biochemical Corporation, CAT# LS003571), and 4.8 mg/mL HEPES in 5 mL HBSS buffer for 60 minutes at 37°C with regular agitation (every 60 s) and the resulting cell suspension was passed through a 70 µm cell strainer again.

To separate surface-adherent from tissue-resident cells, samples were repeatedly agitated in PBS after removal of residual blood until no cells were recovered in the supernatant prior to enzymatic digestion.

### Flow cytometry and cell sorting

Cells were resuspended in a defined volume of staining buffer and incubated with an unconjugated monoclonal antibody (mAb) against CD16/CD32 (eBioscience) to block nonspecific Fc receptor-mediated binding of staining antibodies. Cells were then labelled with fluorochrome-conjugated antibodies for 30 minutes at 4°C. Antibodies used in myeloid cell cytometry and sorting (all from BioLegend, San Diego, CA): CD45-FITC, CD14-BV421, CD68-BV711, HLA-DR-APC, CD206-PECy7, CD11b-BV650, CD163-PE. To gate out lineage cells in Monocyte/macrophage phenotyping PeCy7-conjugated CD3, CD15, CD20, CD56 antibodies were combined.

Antibodies used in T cell flow cytometry (from BioLegend, San Diego, CA or BD Biosciences San Jose, CA): CD3-APC, CD45RA-FITC, CD62L-PerCP Cy5.5, CD4-APC Cy7, CD8-eFluor 450, CD69-PE. IFN-γ-BV711, TNF-α-BV750, G-CSF-APC, GM-CSF-AF647, IL-22-PE, IL-4-BV421, IL-6-PerCP-eFlour710, IL-10-APC-R700, TGF-β-PE-CF594, IL-17A-BV605, CCL4-PE-Cy7. Dead cells were excluded using Fixable Viability Dye eFluor 510 or Zombie NIR. Flow cytometry was performed on a BD LSR II (Becton Dickinson, Heidelberg, Germany) or Cytek Aurora (Cytek Biosciences), and data were analyzed with FlowJo v11.

CD8^+^ and CD4^+^ T_EM_ cells were pre-enriched from leukocyte reduction cone PBMC using the CD4^+^ Effector Memory T Cell Isolation Kit (Miltenyi Biotec) or the MojoSort Human CD8 Memory T Cell Isolation Kit and TCR^+^CD62L^neg^CD45RA^neg^CD56^neg^ cells isolated on a CytoFlex SRT Cell Sorter (Beckman Coulter), achieving a purity of >95%.

### RNA sequencing

Full-length complementary DNA (cDNA) was prepared using 2 ng total RNA (RNA integrity number ˃8) isolated from 300 sorted CD45^+^CD14^+^CD206^neg^ and 300 CD45^+^CD14^+^CD206^pos^ cells using the Smart-Seq V4 Ultra low Input RNA kit (Takarabio) with 12 cycles of final amplification. Quantity was assessed by Invitrogen’s Qubit HS assay, and the size of full-length cDNA was determined using Agilent’s 2100 Bioanalyzer HS DNA chip. Barcoded sequencing libraries were prepared using Illumina’s Nextera XT kit using 1ng cDNA. Barcoded sequencing libraries were onboard clustered using HiSeq Rapid SR Cluster Kit v2 using 8 pM and 59 basepairs and sequenced on the Illumina HiSeq2500 using HiSeq Rapid SBS Kit v2 (59 Cycle). Raw output HiSeq data were preprocessed according to the Illumina standard protocol. Sequence reads were trimmed for adapter sequences and further processed using the software CLC Genomics workbench (v12.0 with CLC’s default settings for RNA-Seq analysis). Reads were aligned to GRCh38 genome.

The bulk RNA-seq data from this project have been deposited in the Gene Expression Omnibus database (GEO)(accession number GSE331428).

### Histology

Vein samples were stored in 15 mL Falcon tubes with approx. 2-3 mL zinc formalin for 24 h. After 24 h, zinc formalin solution was removed and replaced with 70% ethanol.

Deparaffinized sections (5 μm) were immunohistochemically stained using antibodies against CD68 (rabbit polyclonal; abcam) and CD3 (rabbot polyclonal, abcam). Slides were labeled with the primary antibody during incubation at 4 °C overnight. After washing in PBS, sections were incubated in biotinylated secondary antibody for 1 hour (Histofine universal alkaline phosphatase polymer), followed by washing in PBS. Finally, the reaction was visualized by incubation for 15 minutes in alkaline phosphatase (AP-Vectastain, Vector Laboratories, Burlingame, CA). Isotype controls were used accordingly. Slides were analyzed on a slide scanner (NanoZoomer S360 MD, Hamamatsu) and NDP.2 view Software. Positively stained cells on the slide were counted by two independent investigators in a blinded fashion.

### Statistical analyses

Quantitative data are expressed as mean ± standard error of the mean (SEM). Normal data distribution was checked with the Shapiro-Wilk test. Differences between two groups were assessed using an unpaired Student’s *t-*test for normally distributed data, or the Mann– Whitney test for non-normally distributed data. Comparisons among multiple groups were performed using one-way analysis of variance (ANOVA). Statistical significance was defined as a *P* value ˂0.05. All analyses were performed using GraphPad Prism software (version 9; GraphPad Software).

## Results

### Macrophages in the connective tissue and the vein wall of varicose veins differ in CD206 expression

Varicose veins from patients undergoing vein stripping for primary varicosity, as well as non-varicose great saphenous veins harvested for bypasses, were enzymatically digested using a previously developed method that preserves cell surface markers^6^ and the percentage of immune cells determined by flow cytometry. For quantitative comparisons, all specimens underwent identical dissociation and staining workflows.

The proportion of immune cells in varicose vein samples ranged from 0.6 to 8% (mean 3 %, SD 2.2) (**Figure 1A**, **Supplementary Figure S1**). CD45^+^ immune cells in varicose veins contained a high proportion of CD14^+^ cells expressing HLA-DR, CD68 and CD206 (**Figure 1B**) and immunohistochemical staining identified CD68^+^ cells localized in the tunica externa (connective tissue) and tunica media, but not in the tunica intima (**Figure 1C**). In comparison, great saphenous veins contained on average eightfold fewer immune cells (mean 0.26%, SD 0.57, p-value bypass vs. varices ˂0.0001) (**Supplementary Figure S1B)** including very few CD68^+^ cells, which were consistently restricted to the tunica media (**Supplementary Figure S1A**). Due to the very low number of immune cells, detailed flow cytometric analyses could not be performed on great saphenous vein samples.

**Figure 1:**
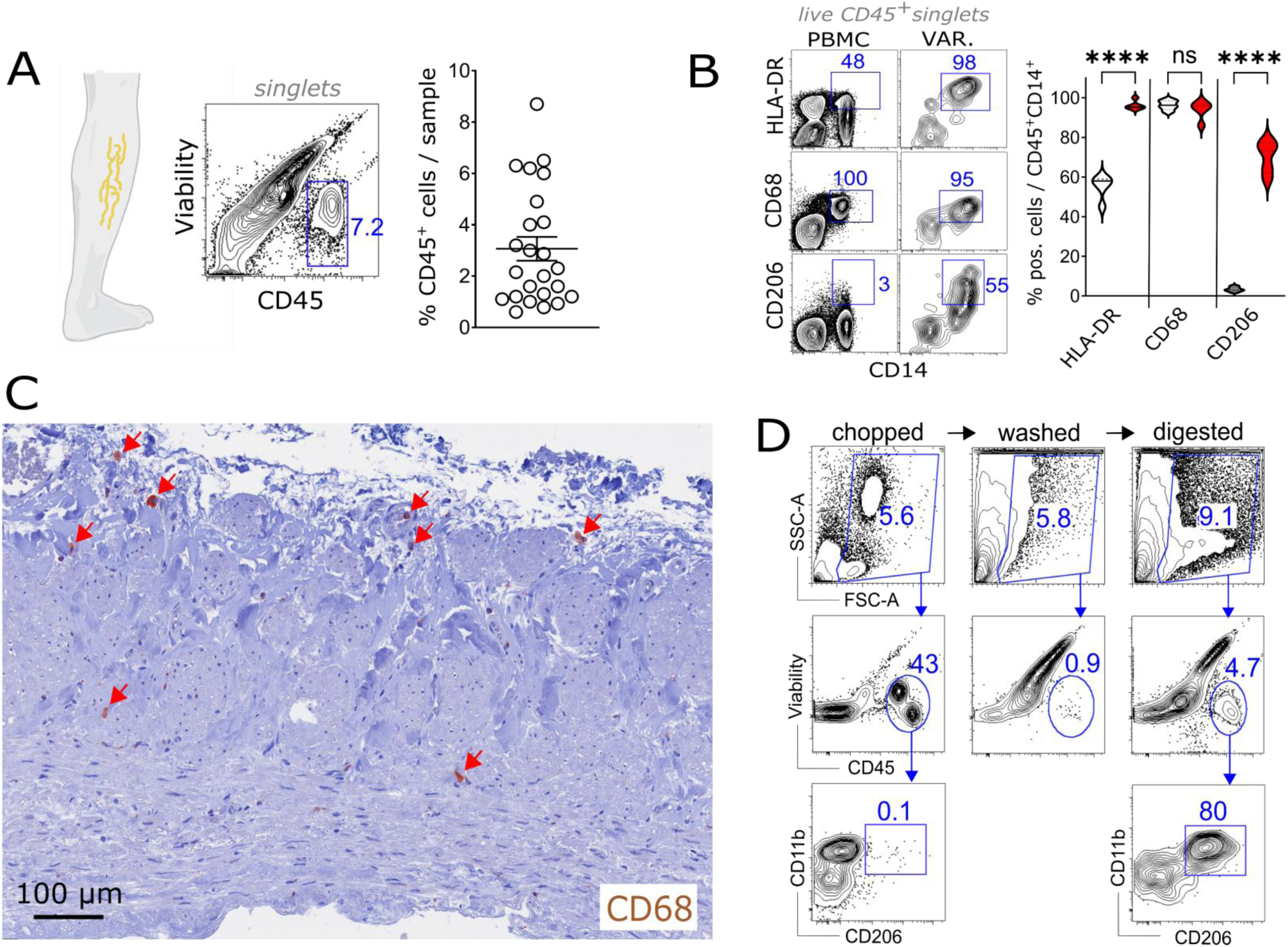
Immune cells in varicose veins. **A)** Flow cytometric analysis of CD45^+^ immune cells in in single cell suspensions of varicose veins (mean 3%, SD 2.2). **B)** Representative flow cytometric analysis of myeloid cell markers in single cell suspensions of varicose tissue and peripheral blood. Violin plots show median, interquartile range, minimum and maximum, n = 5 per sample. **C)** Immuno-histochemical CD68 staining of a varicose saphenous vein FFPE section (×200 magnification, monoclonal anti-CD68 antibody + hematoxylin and eosin staining). **D)** Representative flow cytometric analysis of myeloid cells obtained from varicose veins by mechanical cleaning and subsequent tissue digestion. All data representative of at least five independent experiments, numbers indicate cell frequencies.

Attempts to analyze mRNA expression in CD68^+^ cells across different varicose vein tissue regions by Digital Spatial Profiling (NanoString) were unsuccessful, owing to variable CD45^+^ cell numbers across regions of interest (ROI) and insufficient read counts from the low abundance of CD68^+^ cells relative to total tissue cells.

To identify markers that distinguish immune cells in different tissue regions, varicose veins were subjected to a multi-stage mechanical cleaning process, and the phenotype of the cells obtained in each step was analyzed by flow cytometry. To this end, the veins were first opened longitudinally and rinsed to remove non-adherent blood cells. The tissue was then repeatedly agitated in culture medium to dislodge loosely adherent cells. Next, the tissue was chopped into small pieces (> 1 mm) and subjected to three additional rounds of agitation (washing) in culture medium. Once no further cells could be mechanically separated from the tissue fragments, the remaining tissue was enzymatically digested to release cells from deeper tissue layers (**Figure 1D**). Comparative analysis of separated ‘outside’ (mechanically detached) and ‘inside’ (digestion liberated) CD45^+^CD11b^+^ cells identified CD206 expression as a distinguishing marker (**Figure 1D and Supplemental Figure S2A**). Other markers such as CD163 or CD101 could not discriminate between ‘outside’ and ‘inside’ CD45^+^CD11b^+^ populations. Neither ‘outside’ nor ‘inside’ CD45^+^CD11b^+^ cells expressed granulocyte markers (CD15, CD66b, not shown).

### The connective tissue contains inflammatory active monocytes while tissue-regenerating macrophages develop in the vein wall

To determine the gene expression of loosely attached CD206^neg^ ‘outside’ and CD206^pos^ ‘inside’ monocytic cells, 300 CD45^+^CD14^+^CD206^neg^ and CD45^+^CD14^+^CD206^pos^ cells were sorted from each of five individual varicose vein samples upon tissue digestion without mechanical pre-cleaning (**Figure 2A**). Principal component analysis (PCA, fold change >2, Difference ≥4) (**Figure 2B**) and hierarchical clustering (**Figure 2C**) demonstrated segregation by origin and numerous differences between CD206^pos^ ‘inside’ and CD206^neg^ ‘outside’ myeloid cell gene expression profiles. All preparations confirmed the sorting signature (*CD206*^+/-^) (**Figure 2D**) as well as a shared monocytic phenotype characterized by *CD11b*^+^, *CD33*^+^, *CX3CR1*^+^, *CD114*^+^, and *HLADR*^+^ expression (**Figure 2E**). Consistent with flow cytometric profiling, neither ‘inside’ nor ‘outside’ cells expressed granulocyte markers such as *FUT4* (CD15) or *CEACAM 8* (CD66b) (not shown, dataset publicly available via Gene Expression Omnibus, accession number GSE331428).

**Figure 2:**
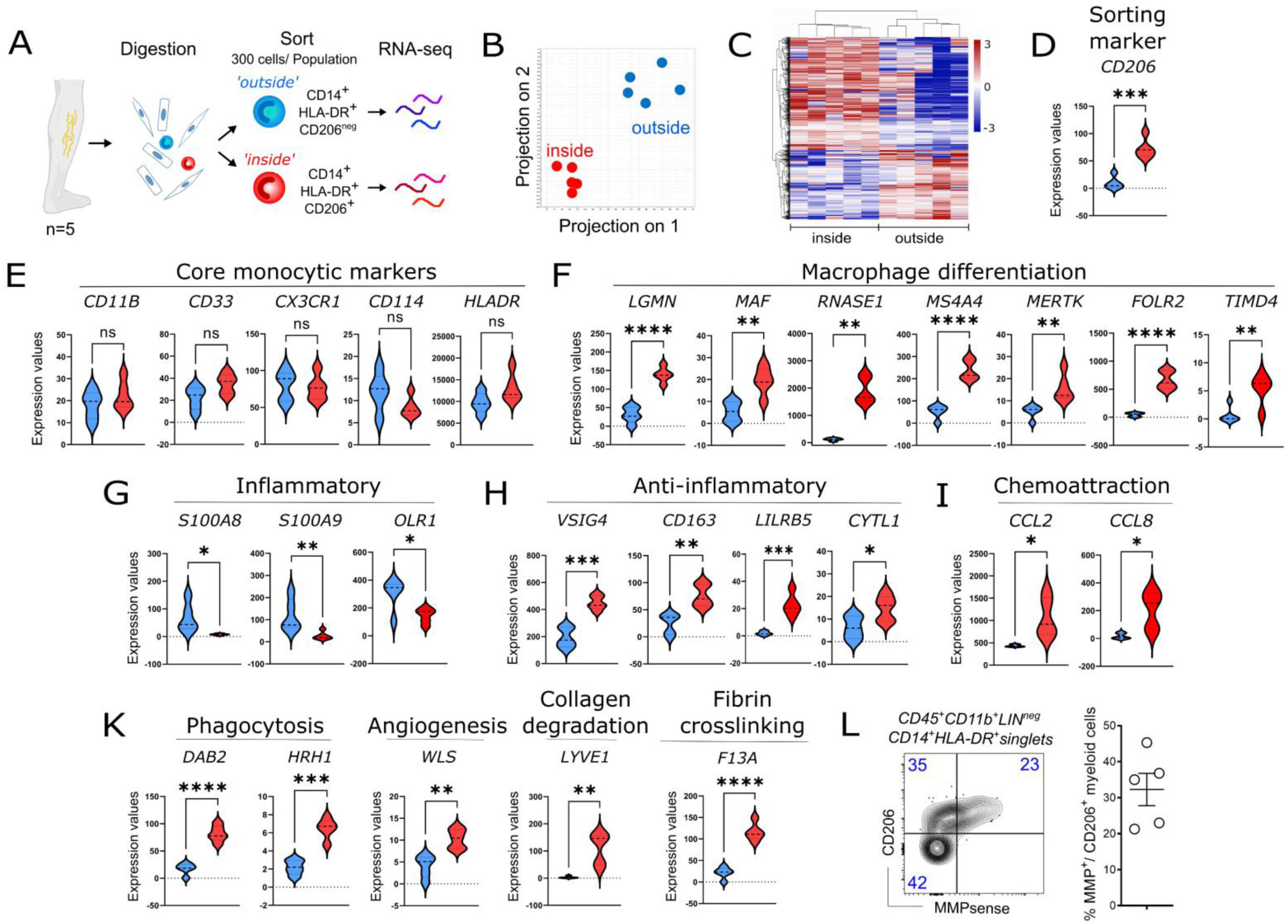
RNA profiles of CD206^neg^ ‘outside’ and CD206^pos^ ‘inside’ varicose monocytic cells. **A)** RNA sequencing experimental workflow (n = 5 independent preparations). **B)** Principal component analysis of RNA signatures in flow-sorted CD14^+^HLADR^+^CD206^neg^ (outside-blue) and CD14^+^HLADR^+^CD206^pos^ (inside-red) populations. **C)** Heat map of regulated genes (fold change >2, Difference ≥4). **D)-K)** Violin plots of reads per kilobase million (RPKM) values for the sort marker CD206 (**D**), core monocytic markers (**E**), macrophage differentiation marker (**F**), inflammatory genes (**G**), anti-inflammatory genes (**H**), Chemokines (**I**) and Phagocytosis, Angiogenesis, Collagen degradation and Fibrin crosslinking-related genes (**K**) in sorted CD14^+^HLADR^+^CD206^neg^ and CD14^+^HLADR^+^CD206^pos^ varicose cells. **L)** MMP activity. Samples were pre-incubated with 4 nmol PBS-reconstituted MMP Sense 645 (Perkin Elmer, Boston, USA) for 24 hours prior to digestion, surface staining and flow cytometric analysis. Left: representative flow cytometric plot, numbers indicate cell frequencies. Numbers indicate percentages of gated live cells; symbols represent individual specimen. Results shown as mean ± SEM.

Among the genes most strongly up-regulated in ‘inside’ CD206^pos^ compared to ‘outside’ CD206^neg^ cells were *LGMN* (Legumain)^7^, *MAF* (Transcription factor Maf),^8^ *RNASE1 (Ribonuclease A family member 1),*^9^ *MS4A4* (Membrane-Spanning 4-Domains, Subfamily A, Member 4),^10^ *MERTK* (Proto-oncogene tyrosine-protein kinase MER),^11^ *FOLR2* (Folate receptor beta)^12^ and *TIMD4*^13^ all of which are associated with resident, reparative, fetal-derived macrophages (**Figure 2F**).

While the ‘outside’ population expressed inflammatory genes *S100A8* and *S100A9* (S100 calcium-binding protein A8 and A9) and *OLR1* (Oxidized Low Density Lipoprotein Receptor 1) (**Figure 2G**) consistent with recently recruited inflammatory monocyte-derived macrophages, the ‘inside’ CD206^pos^ cluster exhibited higher expression of anti-inflammatory genes *vsig4* (V-Set And Immunoglobulin Domain Containing 4),^14^ *CD163* (Scavenger Receptor Cysteine-Rich Type 1 Protein M130),^15^ *LILRB5* (leukocyte immunoglobulin like receptor B5, CD85C), an inhibitory receptor that regulates the antigen-presenting function of cells through intracellular binding of HLA-class I heavy chains,^16^ suggesting reduced antigen-presenting capacity in macrophages within the vein wall. In addition, *CYTL1* (Cytokine Like 1) a regulatory cytokine-like factor was also enriched (**Figure 2H**). The ‘inside’ population further showed an up-regulation of *CCL2* (C-C Motif Chemokine Ligand 2), *CCL8* (C-C Motif Chemokine Ligand 8) and *CYTL1* (Cytokine Like 1) (**Figure 2I**) involved in the recruitment and activation of immune cells, potentially contributing to varicose immune cell infiltration.

In addition, *DAB2* (Disabled homolog 2 Adaptor Protein 2) which regulates macrophage polarization,^17^ *HRH1* (Histamine Receptor H1)^18^ required in phagocytosis, *WLS* (Wnt Ligand Secretion Mediator) involved in angiogenesis,^19^ *LYVE1* (Lymphatic vessel endothelial hyaluronan receptor 1) participating in collagen degradation^20^ and *F13A* (Factor XIII subunit A of blood coagulation) associated with macrophage polarization and Fibrin cross-linking^21^ were upregulated in the ‘inside’ population (**Figure 2K**).

The high degree of agreement among the mRNA data from the patient samples, as well as the strong agreement with the flow cytometry data, underscore the reliability and reproducibility of our results.

Incubation of varicose vein samples with a matrix metalloproteinase (MMP)-sensitive probe prior to tissue digestion revealed that MMP activity (MMP*-*2, MMP*-*3, MMP*-*9, and MMP*-*13) was exclusively restricted to the ‘inside’ vein wall CD206^pos^ population (**Figure 2L**).

Together, these gene expression profiles indicate the presence of active inflammatory, likely monocyte-derived (injury-responsive) macrophages in the connective tissue, alongside activated, tissue-resident, remodeling macrophages in the vein wall.

### Varicose vein macrophage cytokines induce T cell IFN-γ production *in situ*

In addition to monocytic cells, varicose veins have been reported to contain T cells^4, 6, 22^ and we predominantly identified T cells within the connective tissue (**Figure 3A**). Flow cytometric analysis revealed a primarily effector memory phenotype (CD45RA^neg^ CD62L^neg^)^6^ (**Figure 3B** and **Supplemental Figure S2B**) along with markers of local activation/retention (CD69^+^) **(Figure 3C)**. However, these T cells lacked markers associated with mucosal and barrier tissue residency (CD103, data not shown).

**Figure 3:**
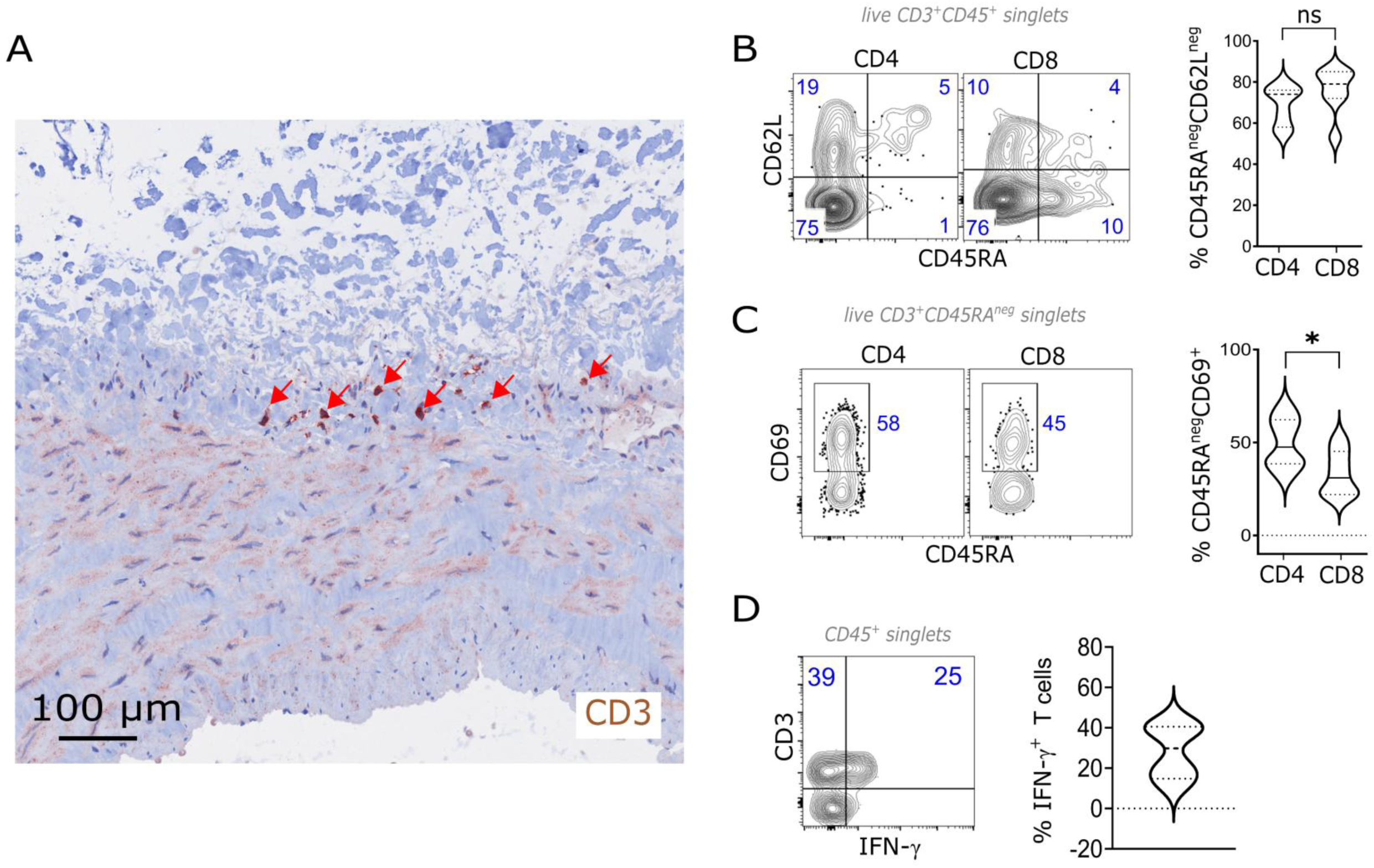
T cells in varicose veins. **A)** Immuno-histochemical CD3 staining of a varicose saphenous vein FFPE section (×200 magnification, monoclonal anti-CD3 antibody + hematoxylin and eosin staining). **B)-D)** Flow cytometric analyses of T cells in varicose veins. **B)** CD45RA and CD62L expression by T cells delineates 4 subsets: naive (T_N_: CD62L^+^CD45RA^+^), central memory (T_CM_: CD62L^+^CD45RA^neg^), (T_EM_: CD62L^−^CD45RA^neg^), and terminal effector (T_EMRA_: CD62L^neg^CD45RA^+^) T cells. **C)** CD69 expression by CD45^neg^ T cells identifies CD69^+^ memory T cells. **D)** Interferon-gamma (IFN-γ) expression in varicose vein T cells. Varicose veins were cultured over night and Brefeldin A added for 1h prior to digestion and flow cytometry. Numbers indicate percentages of gated live cells within the total number; symbols represent individual specimen. Results shown as mean ± SEM.

A previous publication showed that lymphocytes from incompetent great saphenous veins produced Th1 cytokines upon *ex vivo* PHA stimulation.^23^ However, since *ex vivo* stimulation does not necessarily reflect actual T cell behavior *in vivo*, we assessed T cell interferon (IFN) gamma production in varicose veins *in situ.* To this end, varicose vein samples were briefly incubated with brefeldin A prior to tissue digestion and flow cytometric analysis. We detected IFN-γ production by T cells in varicose veins (Figure 3D), with considerable inter-sample variability (**Figure 3D**).

The function of T cells is closely linked to their T cell receptor (TCR)-mediated antigen specificity; however, it is increasingly recognized that T cells lacking disease-relevant TCR specificities can play critical roles in numerous pathological conditions,^24, 25^ including thrombosis.^6, 26^ In particular, memory T cells can be activated to produce cytokines by nonspecific extrinsic signals, especially cytokines, a phenomenon referred to as “bystander” or “innate” activation.^6, 24, 27, 28^ To elucidate the mechanisms driving T cell activation and IFN-γ production in varicose vein tissue, we revisited the macrophage gene expression dataset derived from these samples.

The mRNA expression data of monocytic cells in varicose veins revealed the expression of cytokine genes *IL1A, IL1B, IL6, IL15, IL16, IL18, IL23A* and *TNFA* (**Figure 4A**). With the exception of *IL1a*, which was more highly expressed in the ‘outside’ population and *IL23a*, no significant differences were observed between the ‘outside’ and ‘inside’ populations. To determine which varicose-vein derived monocytic cytokines stimulate T_EM_ cells (the predominant infiltrating T cell subtype), and which factors are subsequently produced by T_EM_ in response, isolated blood-derived CD8^+^ and CD4^+^ T_EM_ cells were stimulated with a recombinant cytokine cocktail reflecting the monocytic cytokine profile, as well as with “minus-one” cocktail variants lacking individual components.

**Figure 4:**
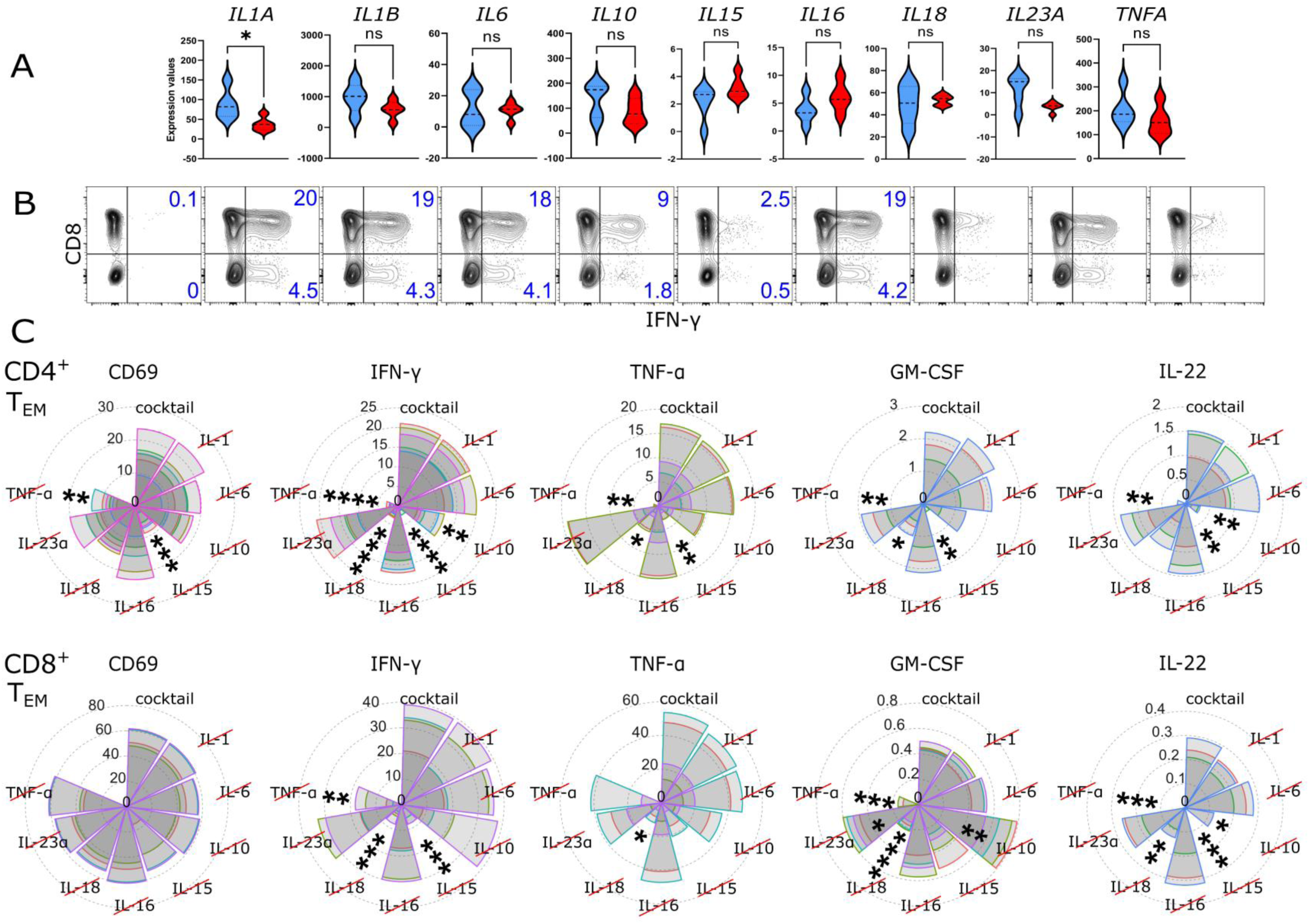
Effector-memory T cells (T_EM_) are activated by cytokines expressed in varicose vein monocytes. **A)** Cytokine gene expression of CD206^neg^ ‘outside’ (blue) and CD206^pos^ ‘inside’ (red) varicose monocytic cells. **B)** IFN-γ production by sorted CD8^+^ T_EM_ 24 h after stimulation with varicose monocyte cytokines cocktail (upregulated cytokines shown in (**A**) or minus-1 variants (2 ng/ml each)). Representative flow cytometric plots. **C)** CD69 up-regulation, IFN-γ, TNF-α, IL-22 and GM-CSF production by flow-sorted CD8^+^ and CD4^+^ T_EM_ 24 h after stimulation with varicose monocyte cytokines cocktail variants. Cumulative data representative of 4 individual experiments. All results shown as mean ± SEM. Comparisons made to full cocktail stimulation.

The complete cytokine cocktail induced robust IFN-γ production in both CD8⁺ and CD4⁺ T_EM_ cells (**Figure 4B**). Slightly fewer cells produced TNF-α, and only a small fraction of T_EM_ cells produced IL-22 and GM-CSF (**Supplemental Figure S3 and S4**). Additional analyses excluded the production of IL-4, IL-6, IL-10, IL-17A or TGF-β. “Minus-one” variants of the cytokine cocktail confirmed the previous reported key roles of IL-15 and IL-18 in driving IFN-γ production.^29, 30^ Notably, IL-10 and TNF-α also significantly influenced the frequency of IFN-γ-producing cells, whereas omission of other cytokines from the cocktail had no detectable effect (**Figure 4B and Supplemental Figure S3)**. The relative importance of stimulatory cytokines (IL15 > IL18 > TNF > IL-10) appeared consistent across all induced cytokines.

## Discussion

Varicosis of the veins is increasingly recognized as a chronic inflammatory and progressively degenerative vascular disease in which immune-mediated mechanisms act alongside hemodynamic dysfunction. Although leukocyte infiltration and elevated inflammatory mediators have been documented in varicose tissue for more than two decades,^3–5^ the functional identity and spatial organization of the dominant myeloid cell populations have remained poorly defined. By combining stepwise mechanical tissue compartmentalization with transcriptomic profiling of flow-sorted myeloid cells, and by directly interrogating the functional impact on infiltrating T cells, we provide evidence that varicose veins harbor two spatially and functionally distinct macrophage populations that sustain chronic venous inflammation through complementary and mutually reinforcing mechanisms.

The perivascular connective tissue harbored CD206^neg^ IL-1α^+^ S100A8/9^+^ macrophages characterized by a strongly pro-inflammatory transcriptomic signature, whereas the vein wall contained CD206^pos^ TIMD4^+^ LYVE1^+^ macrophages associated with tissue-remodeling and homeostatic functions. This anatomical dichotomy of macrophage identity has not previously been described in the venous system and parallels the two interstitial macrophage niches reported across non-lymphoid organs by Chakarov et al. (nerve-associated LYVE1^lo^MHC-II^hi^ vs. vessel-associated LYVE1^hi^MHC-II^lo^, the latter homeostatic).^31^ Vein wall macrophages share the LYVE1^hi^, anti-inflammatory gene signature and are described here for the first time in venous tissue. Whether they are embryonically seeded or postnatally recruited and locally imprinted cannot be resolved transcriptomically; however, strong expression of TIMD4, FOLR2, MAF, MERTK, and LGMN, canonical markers of fetal-liver-derived, long-lived tissue-resident macrophages^11–13^ supports a resident origin. MMP activity was exclusive to this inner population, identifying it, despite an otherwise homeostatic profile, as the principal driver of collagen turnover and structural wall remodeling.

Connective-tissue macrophages instead expressed S100A8, S100A9, and OLR1 (LOX-1), a profile typical of recently recruited, monocyte-derived macrophages responding to venous hypertension.^32^ S100A8/9 are alarmins that, via TLR4/RAGE, drive NF-κB signaling and leukocyte recruitment.^33^ This fits a model in which venous hypertension increases endothelial permeability, allowing blood-derived signals into the perivascular space and amplifying S100A8/9-driven innate inflammation. Co-expression of OLR1 further suggests oxidative/metabolic stress, as in atherosclerosis. Previously reported elevations of IL-1β and TNF-α,^34^ can now be localized specifically to this connective-tissue macrophage compartment, a spatial resolution not achievable by prior immunohistochemistry or bulk analyses.

Both macrophage populations expressed IL1A, IL1B, IL6, and TNFA, indicating that chronic inflammation is not restricted to one subtype, consistent with a functional continuum rather than a strict M1/M2 dichotomy.^11^ Vein wall macrophages, despite expressing anti-inflammatory markers (VSIG4, CD163, LILRB5, CYTL1), continued producing inflammatory cytokines - a state resembling “frustrated resolution” seen in rheumatoid arthritis and atherosclerosis.^35^ Upregulation of DAB2 (polarization regulator) and WLS (Wnt-dependent angiogenesis) further implicates these cells in abnormal vascular remodeling. IRF1, recently proposed as required for monocyte-to-macrophage differentiation *in vitro*,^36^ was not differentially expressed, leaving the differentiation trajectory unresolved pending single-cell/lineage-tracing approaches.

Elevated CCL2, previously been reported in varicose tissue,^37^ and CCL8 in vein wall macrophages suggest active recruitment of monocytes and T cells. We now identify LYVE1^+^ vein wall macrophages as a major source of CCL2, implying a self-reinforcing recruitment loop that could sustain inflammation independent of the initial hemodynamic trigger. F13A and MMP family genes, highly expressed here, have previously been linked to poor outcomes in non-healing venous ulcers,^38, 39^ suggesting a molecular continuum toward more severe chronic venous insufficiency (CVI).

T cells were predominantly effector-memory (T_EM_), CD69^+^ tissue-resident and/or activated, and localized in the perivenous connective tissue. Brefeldin A trapping demonstrated active IFN-γ production *in situ*. Several findings argue against classical TCR-driven activation: no auto-antigen meeting Witebsky’s criteria has been identified,^40^ costimulatory molecules (CD86, CD40, CD54) were not differentially expressed; both macrophage populations expressed IL16, a suppressor of TCR-mediated activation^41^ and LILRB5 was upregulated in vein wall macrophages. This supports TCR-independent bystander activation of resident memory T cells^24, 25, 42^ by a cytokine milieu (IL-1α, IL-1β, IL-6, IL-15, IL-16, IL-18, IL-23a, TNF-α) sufficient to induce IFN-γ in isolated CD4^+^/CD8^+^ T_EM_ cells without TCR stimulation. IL-15 and IL-18 were dominant drivers, consistent with STAT5-mediated IL-18R upregulation and GADD45β-dependent IFN-γ induction.^29^ TNF-α acted co-stimulatory;^29^ unexpectedly, IL-10 also enhanced responsiveness, likely via STAT3-mediated IL-2 receptor upregulation. CD4^+^ T_EM_ cells depended on all four cytokines (IL-15, IL-18, TNF-α, IL-10), while CD8^+^ T_EM_ cells showed greater redundancy.^43, 44^ Given that both subsets increase with age, and that we previously showed age-dependent bystander activation in a murine thrombosis model,^6^ this mechanism may contribute to age-related CVI progression.

T cell-derived IFN-γ likely further activates macrophages (IL-1β, TNF-α, MMP expression), particularly within the CD206^neg^ population, a circuit described in psoriasis, sarcoidosis, and atherosclerosis, and now mechanistically defined for CVI, extending earlier histological co-localization findings.^4, 22^ Confinement of immune cells to the tunica externa/media (while absent from the intima), together with clear transcriptomic differences from circulating monocytes, supports entry via the vasa vasorum rather than the lumen, consistent with venous hypertension-driven adventitial remodeling and vasa vasorum expansion in CVI.^45^

Several aspects of our study warrant methodological consideration. Mechanical pre-cleaning identified two macrophage populations and CD206 proved to be the key discriminating marker empirically (CD163 and CD101 were not equivalent). Smart-seq V4 on 300 sorted cells yielded sufficient depth despite limited numbers; consistency across five patients and concordance with flow cytometry support robustness. The lower frequency of IFN-γ^+^ T cells *in situ* versus after *ex vivo* stimulation likely reflects the low-grade inflammation of C2-stage disease rather than a technical limitation. It should also be noted that vein segments obtained during surgical procedures, including those harvested for bypass grafting, do not represent truly healthy control tissue, as even macroscopically unaffected venous segments may already harbor subclinical inflammatory or remodeling changes. We also acknowledge that protein abundance, spatial source, and IL-15 trans-presentation were not assessed *in situ*.

## What are the Clinical Implications?

Current treatment of varicose veins remains largely hemodynamic-compression, sclerotherapy, ablation, or surgery-and does not address the underlying inflammatory drivers of disease. By identifying two functionally distinct macrophage populations that jointly sustain a self-reinforcing myeloid-lymphoid inflammatory circuit, this work defines a previously unrecognized cellular and molecular framework for chronic venous disease that is, in principle, amenable to targeted immunomodulation. The demonstration that macrophage-derived IL-15, IL-18, TNF-α, and IL-10 drive TCR-independent bystander activation of effector-memory T cells suggests that interrupting this cytokine axis could dampen chronic inflammation while preserving protective immunity. Similarly, the S100A8/9-TLR4/RAGE amplification loop in connective-tissue macrophages and the MMP-driven remodeling activity of vein wall macrophages represent compartment-specific therapeutic targets, potentially allowing intervention tailored to disease stage. Given that IL-1 pathway blockade has already shown efficacy in atherosclerosis,^46^ an analogous anti-inflammatory strategy may be feasible in CVI, potentially slowing progression from uncomplicated varicose veins to more severe manifestations such as venous ulceration. Longitudinal and interventional studies will be needed to determine whether targeting this macrophage-T cell axis can reduce disease progression, recurrence after intervention, or ulcer formation, and whether specific cytokine signatures could serve as biomarkers to stratify patients for future immunomodulatory therapies.

